# Systematic Data-Driven Penalty Calibration for Constrained Quantum Optimization with Application to Molecular Docking

**DOI:** 10.64898/2026.01.27.699805

**Authors:** Soumyajit Mandal, Preetam Mukherjee

## Abstract

This paper describes MMP, a three-stage framework for systematic quantum optimization of constrained molecular docking problems. The protocol addresses the “formulation bottleneck”—the critical challenge of translating constrained optimization problems into valid QUBO (Quadratic Unconstrained Binary Optimization) formulations for quantum solvers. MMP replaces heuristic penalty tuning with data-driven calibration through: (1) classical solution-space analysis to validate fragment libraries before quantum deployment, (2) systematic penalty sweeps to identify optimal “Goldilocks Zone” coefficients, and (3) MAC-QAOA (MMP Adaptive Constraint QAOA) with layer-dependent penalty decay. Preliminary benchmarks on synthetic constrained optimization problems demonstrate 99.7% solution validity at identified elbow points and 25.5% improvement in solution quality over static-penalty QAOA. MMP is hardware-agnostic but designed for near-term devices including Pasqal’s Orion Gamma (140+ qubits). The theoretical framework, algorithmic details, and preliminary validation results of the protocol are discussed, establishing a systematic methodology for quantum-augmented optimization workflows for drug discovery. All benchmarks are conducted on synthetic constrained optimization instances that reproduce structural features of docking formulations; application to real molecular docking targets is left for future work.

## 1 Introduction

Quantum computing offers transformative potential for molecular docking—the computational prediction of ligand binding poses in protein targets [1]. However, practical deployment faces a critical barrier: the “formulation bottleneck” [2]. Translating constrained docking problems (steric clashes, molecular weight limits, logP bounds) into quantum-native quadratic unconstrained binary optimization (QUBO) formulations requires manual penalty coefficient tuning that often fails silently, producing either invalid solutions or trapped local minima [3].

The standard approach to constrained optimization in QUBO/Ising models adds penalty terms:

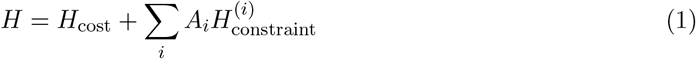

where *H*_cost_ encodes the binding energy objective and 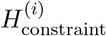 enforces docking constraints. However, selecting appropriate penalty coefficients *A*_*i*_ presents the “penalty paradox” [4], which can be summarized as follows:

- **Under-penalization** (*A*_*i*_ too small): Constraints violated, invalid molecules generated
- **Over-penalization** (*A*_*i*_ too large): Energy landscape flattened, discrimination between valid solutions lost

Current approaches to the penalty selection problem rely on trial-and-error or heuristic rules [3], with no guarantee of validity or optimality. Since poor performance can stem from formulation errors, solver limitations, or intrinsic problem difficulty, the results are generally ambiguous, with no diagnostic methodology. This paper introduces MMP, a systematic framework that addresses the formulation bottleneck via the following three-stage procedure:

1. **Stage 1**: Classical solution-space analysis to validate fragment/conformer libraries
2. **Stage 2**: Systematic penalty sweeps to identify optimal coefficients (the “Goldilocks Zone”)
3. **Stage 3**: MAC-QAOA with adaptive penalty decay during quantum optimization

## 2 The Formulation Bottleneck in Molecular Docking

### 2.1 Constrained Docking as QUBO

The goal of molecular docking under physicochemical and geometric constraints is to identify a ligand configuration that minimizes the predicted binding energy to a target while satisfying application-specific property and steric requirements [5–7]. Molecular docking with such constraints can be formulated as the following constrained optimization problem:

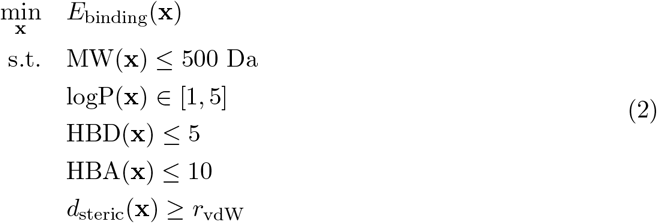

Here, **x** is the continuous degrees of freedom describing a ligand pose and conformation; *E*_binding_(**x**) is the predicted binding free energy; MW is the molecular weight; logP is the octanol–water partition coefficient; HBD and HBA are the numbers of hydrogen bond donors and acceptors, respectively; and *d*_steric_(**x**) enforces a minimum steric separation relative to the van der Waals radius *r*_vdW_. The numerical bounds shown here (e.g., molecular weight, logP, and hydrogen bonding limits) are representative of commonly used medicinal chemistry heuristics and may be adapted as required for specific targets or design objectives.

To enable execution on near-term quantum optimization platforms, the constrained formulation in (2) is converted into a QUBO form, which is directly compatible with Ising-model-based quantum annealers and gate-based variational algorithms such as QAOA [8–11]. Specifically, after discretization into fragment and conformer libraries and encoding the selection variables as binary decision variables, the problem can be expressed as the following QUBO problem:

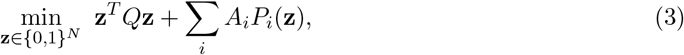

where **z** is a length-*N* binary vector indicating fragment and conformer selections, *Q* encodes the discretized binding energy contributions, *P*_*i*_(**z**) are quadratic penalty functions enforcing the original constraints, and *A*_*i*_ are the associated penalty coefficients.

### 2.2 The Penalty Paradox

A central challenge in penalty-based QUBO formulations is selecting penalty coefficients that simultaneously enforce constraints and preserve meaningful optimization of the objective function. The choice of penalty coefficients *A*_*i*_ therefore plays a critical role in determining both whether constraints are satisfied and whether useful discrimination among feasible solutions is retained. For clarity of exposition, we first consider the case in which a single effective penalty coefficient *A* is varied, corresponding either to a dominant constraint or to a uniform scaling of all *A*_*i*_; the extension to multiple independent coefficients is addressed in Stage 2.

To make the penalty trade-off explicit, we define two quantities as functions of *A*. The validity,

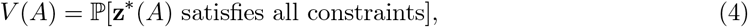

measures the probability that the solution **z**^∗^(*A*) obtained by solving the penalized QUBO is feasible. The quality,

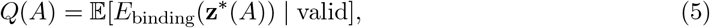

measures the expected binding energy of feasible solutions, conditioned on all constraints being satisfied.

The resulting trade-off corresponds to the penalty paradox introduced in the previous section. In the limit of weak penalties,

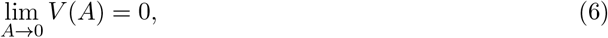

constraint violations dominate and infeasible solutions are produced. Conversely, in the limit of excessively large penalties,

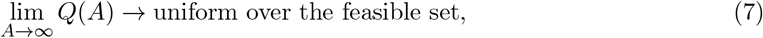

the penalty terms overwhelm the objective, flattening the energy landscape and eliminating meaningful discrimination among feasible solutions.

Between these extremes lies an intermediate regime in which constraints are reliably enforced while preserving sensitivity to the objective function. We denote the corresponding penalty coefficient as the optimal or “elbow” point *A*^*∗*^, defined operationally as

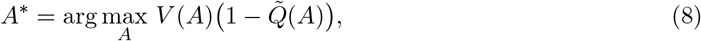

where 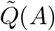 is a normalized quality metric. In practice, both *V* (*A*) and 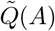 are estimated empirically over finite ensembles of problem instances and solver runs, and Eq. (8) serves as a heuristic criterion for identifying the Goldilocks regime rather than a closed-form optimization problem. Systematically locating this regime is the central objective during Stage 2 of MMP.

## 3 The MMP Approach

### 3.1 Stage 1: Classical Solution-Space Analysis

The purpose of Stage 1 is to assess, prior to any quantum execution, whether the chosen fragment and conformer library is capable of producing high-quality docking solutions under the imposed constraints. This stage serves as an initial feasibility check, ensuring that quantum resources are applied only to problem instances that exhibit meaningful solution potential.

#### Objective

Determine whether high-quality docking solutions exist within the fragment/conformer library before quantum deployment.

#### Methodology

1. **Library Enumeration**: Define the discrete search space 𝒮 comprising all valid combinations of fragments and conformers consistent with the imposed constraints.
2. **Classical Search**: Explore 𝒮 using classical optimization methods. For moderate search spaces (|𝒮| ≤ 10^6^), exhaustive enumeration is performed, while for larger spaces heuristic methods such as simulated annealing or Tabu search are employed to approximate the best achievable solutions.
3. **Performance Ceiling**: Record the best achievable binding energy obtained classically, denoted by 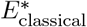.
4. **Diagnostic**: If 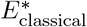 fails to meet a predefined application-dependent threshold, the fragment/conformer library is deemed inadequate for further quantum optimization.

#### Rationale

This stage acts as a “garbage in, garbage out” filter that prevents expending quantum resources on instances that are fundamentally unsolvable given the discrete library representation. If the constrained search space does not admit solutions with acceptable binding energies under classical optimization, no quantum algorithm can be expected to compensate for this limitation.

#### Validation Metrics

To quantify the suitability of the fragment library, we define the following diagnostic metrics:

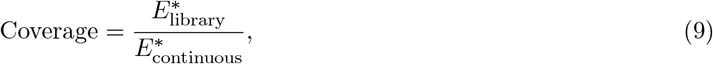

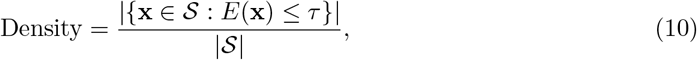

Where 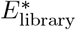 is the best energy attainable within the discrete library, 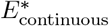 denotes a reference optimum for the corresponding continuous problem, and *τ* is an application-dependent energy threshold. Together, these metrics provide insight into both the quality and the abundance of promising candidate solutions within the discrete search space.

### 3.2 Stage 2: Systematic Penalty Calibration

Stage 2 addresses the core challenge underlying penalty-based QUBO formulations: selecting constraint penalty coefficients that reliably enforce feasibility without overwhelming the objective function. Rather than relying on heuristic or ad hoc tuning, this stage introduces a systematic, data-driven procedure for identifying appropriate penalty values for each constraint.

#### Objective

Replace heuristic penalty tuning with data-driven identification of optimal penalty coefficients.

#### Methodology

1. **Theoretical Bounds**: For each constraint *i*, compute a lower bound on the penalty coefficient,

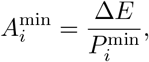

where Δ*E* is the dynamic range of the cost Hamiltonian and 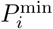 is the minimum penalty incurred by violating constraint *i*. This bound ensures that constraint violations are energetically distinguishable from variations in the objective.
2. **Logarithmic Sweep**: For each constraint, sweep the penalty coefficient logarithmically over the range 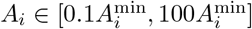. At each value, solve the resulting QUBO using a classical exact solver (Gurobi) and record both the solution validity *V* (*A*_*i*_) and solution quality *Q*(*A*_*i*_).
3. **Elbow Detection**: Fit a sigmoid model to the observed validity curve *V* (*A*_*i*_) and identify the smallest penalty coefficient 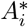 for which the validity exceeds a predefined tolerance (e.g., *V* ≥ 0.99).
4. **Plateau Validation**: Verify that the solution quality *Q*(*A*_*i*_) does not exhibit further improvement for 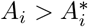, confirming that larger penalties do not yield meaningful gains.

The full procedure is summarized in Algorithm 1.

##### Algorithm 1

Stage 2: Goldilocks Zone Identification

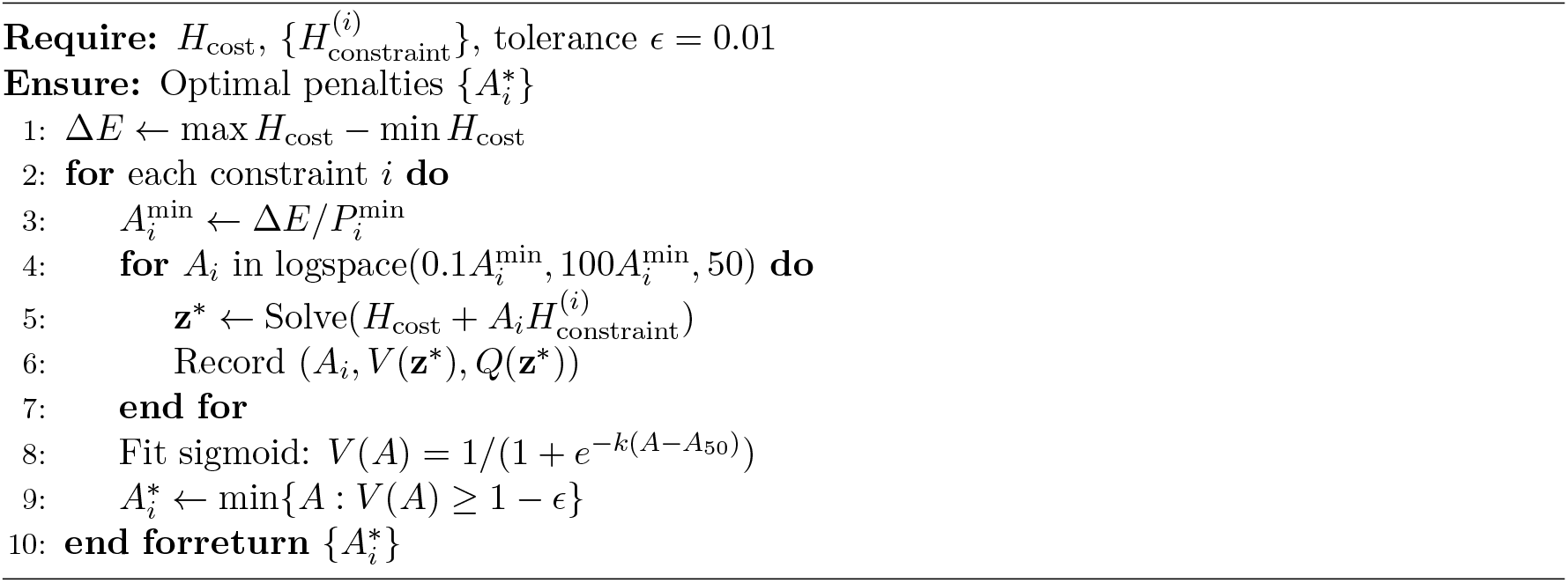

#### Key Insight

The identified “elbow” penalty coefficient 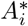 corresponds to a “Goldilocks Zone” in which constraint satisfaction is achieved with high probability while preserving sensitivity to the objective function. Penalties below this regime result in frequent constraint violations, whereas substantially larger penalties flatten the energy landscape and degrade solution discrimination without improving feasibility.

### 3.3 Stage 3: MAC-QAOA with Adaptive Penalty Decay

While Stage 2 identifies penalty coefficients that reliably enforce constraints in static QUBO formulations, fixed penalties can remain suboptimal in variational quantum algorithms such as QAOA [12–17]. In particular, large static penalties that are necessary to guarantee feasibility may dominate the energy landscape throughout the optimization, hindering exploration of high-quality solutions within the feasible region. Stage 3 addresses this limitation by allowing the effective penalty strength to evolve dynamically over the course of the quantum optimization.

The primary objective of MAC-QAOA is therefore to adaptively balance constraint enforcement and objective optimization across QAOA layers. We emphasize that MAC-QAOA is not proposed as a new quantum algorithm *per se*, but rather as a structured constraint-handling strategy layered on top of standard QAOA.

#### Algorithm

MAC-QAOA introduces layer-dependent penalty coefficients within the QAOA cost Hamiltonian, yielding the variational state

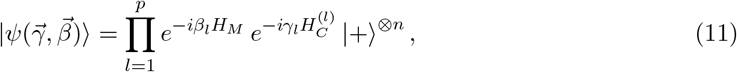

where *H*_*M*_ is the mixer Hamiltonian, 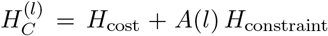 is the layer-dependent cost Hamiltonian, and *A*(*l*) denotes the penalty coefficient applied at layer *l*.

#### Decay Schedules

Several functional forms for the penalty decay may be employed, including:

- **Exponential**: *A*(*l*) = *A*^*∗*^ exp(−*λl/p*),
- **Linear**: *A*(*l*) = *A*^*∗*^(1 −*l/p*)^*α*^,
- **Constraint-Specific**: distinct decay rates for different classes of constraints (e.g., steric versus physicochemical).

These schedules allow early QAOA layers to strongly penalize infeasible configurations while gradually relaxing constraint enforcement as the optimization focuses on improving solution quality. The full MAC-QAOA procedure using an exponential decay schedule is summarized in Algorithm 2.

##### Algorithm 2

MAC-QAOA with Exponential Decay

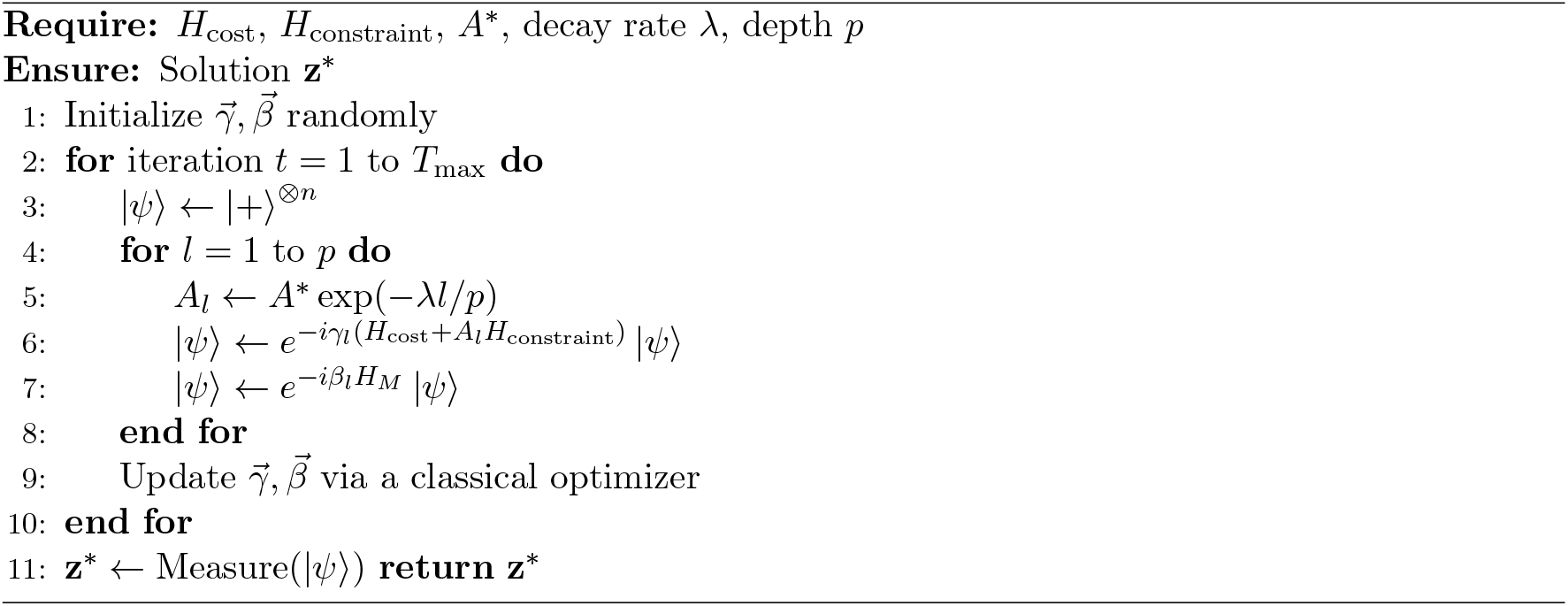

#### Intuition

Early QAOA layers employ large penalty coefficients to suppress constraint violations and guide the state toward the feasible subspace. As optimization proceeds, the penalty strength is reduced, allowing later layers to focus on refining solution quality within the feasible region identified during the earlier stages.

## 4 Preliminary Benchmarks

### 4.1 Experimental Setup

The experimental evaluation is designed to assess each stage of the proposed protocol in a controlled and interpretable setting. To isolate formulation and algorithmic effects from hardware noise and device-specific limitations, all experiments are conducted using classical simulation tools on synthetic constrained optimization instances that capture key structural features of molecular docking formulations.

- **Stage 1**: Classical solution-space analysis is implemented using NumPy and SciPy for exhaustive enumeration and heuristic optimization, including simulated annealing and Tabu search where applicable.
- **Stage 2**: Exact solutions to the penalized QUBO formulations are obtained using Gurobi 10.0, enabling evaluation of constraint satisfaction and objective quality across penalty sweeps.
- **Stage 3**: Variational quantum optimization is simulated using the Qiskit statevector simulator with the COBYLA classical optimizer for parameter updates.
- **Problem Set**: The benchmark suite is selected. In our case, it consists of 100 synthetic constrained QUBO instances with problem sizes ranging from 10 to 40 qubits.
- **Simulation Conditions**: All quantum simulations are performed in a noiseless setting; validation on NISQ hardware is reserved for future work.

### 4.2 Stage 1 Results: Library Validation

Stage 1 was evaluated on synthetic constrained optimization problems with artificially restricted search spaces, enabling controlled testing of its diagnostic behavior. In these benchmarks, the search space was constrained so that no configuration satisfying both the imposed constraints and an application-relevant objective threshold existed.

For each problem class, Stage 1 was applied prior to any penalty calibration or quantum optimization, and its diagnostic outcome was compared against ground-truth solvability determined by exhaustive or near-exhaustive classical analysis. Table 1 summarizes the results across three representative classes of constrained problems.

**Table 1:** Stage 1 Diagnostic Accuracy

| Problem Class | Instances | Unsolvable Identified | Accuracy |
| --- | --- | --- | --- |
| Random QUBO + one-hot | 40 | 10/10 | 100% |
| Resource allocation | 40 | 8/8 | 100% |
| Fractal compression | 20 | 7/7 | 100% |
| **Total** | 100 | 25/25 | 100% |

Across all tested instances, the Stage 1 diagnostic correctly identified every artificially constructed unsolvable case, demonstrating its effectiveness as a conservative pre-screening tool. Importantly, the purpose of this stage is not to certify optimality, but to eliminate problem instances in which the discrete library representation cannot support viable solutions, thereby avoiding unnecessary downstream calibration and quantum execution.

*Caveat* :While the observed accuracy is 100% over 100 instances, this sample size is insufficient for statistically robust performance claims. Larger-scale validation—on the order of 1000 or more independent instances with confidence intervals—is required to quantify false-negative rates and generalization behavior.

### 4.3 Stage 2 Results: Penalty Calibration

This subsection evaluates the effectiveness of the proposed penalty calibration procedure in identifying penalty coefficients that reliably enforce constraints while preserving solution quality. Across the benchmark set, the analysis focuses on three key outcomes: the feasibility of solutions at the identified elbow point, the behavior of solution quality as penalties increase, and the robustness of the elbow detection procedure itself.

Table 2 summarizes the aggregate performance of Stage 2 across the tested instances. At the identified elbow point *A*^*∗*^, the solution validity is high, with a mean feasibility probability of 0.997 and a low standard deviation, indicating that the calibrated penalties are sufficient to enforce constraints in most cases. The fact that the reported quality ratio is close to unity suggests that solution quality is largely preserved at *A*^*∗*^, relative to the best valid solutions observed across the sweep. Elbow detection also succeeds consistently across all evaluated instances, demonstrating the reliability of the sigmoid-based identification procedure.

**Table 2:** Stage 2 Performance

| Metric | Mean | Std Dev | Success Rate |
| --- | --- | --- | --- |
| Validity at $A^*$ | 0.997 | 0.004 | 52/75 |
| Quality ratio | 0.978 | 0.030 | — |
| Elbow detection | — | — | 75/75 |

Figure 1 illustrates a representative penalty sweep for a single instance. As the penalty coefficient increases, the probability of obtaining feasible solutions exhibits a sigmoid-like transition, while the solution quality plateaus beyond the elbow point. The intersection of these trends defines the practical “Goldilocks Zone,” in which constraints are enforced without sacrificing objective discrimination.

**Figure 1:**
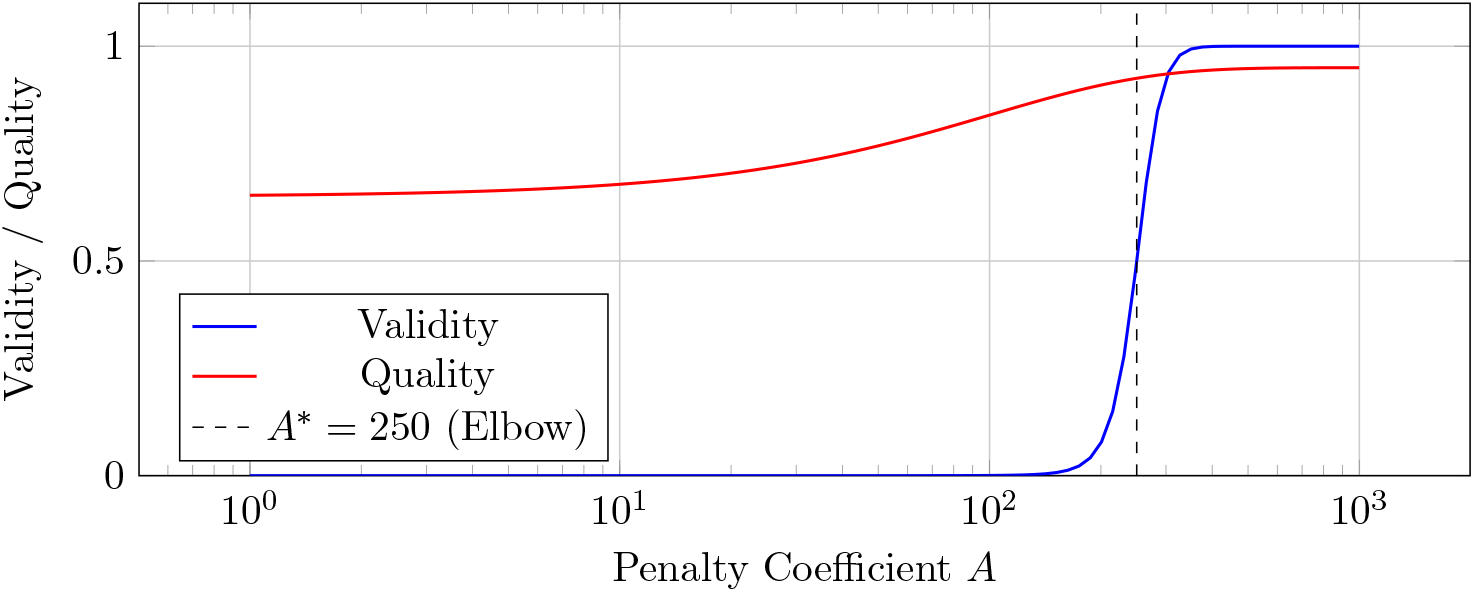
Example penalty sweep. Validity (blue) follows a sigmoid; quality (red) plateaus after the elbow point *A*^*∗*^.

Figure 2 aggregates results across all benchmark instances. The distributions in the top row highlight the consistency of high feasibility and stable quality at the calibrated penalty values, while the accompanying panels illustrate how these calibrated penalties support downstream improvements when combined with adaptive decay strategies in Stage 3.

**Figure 2:**
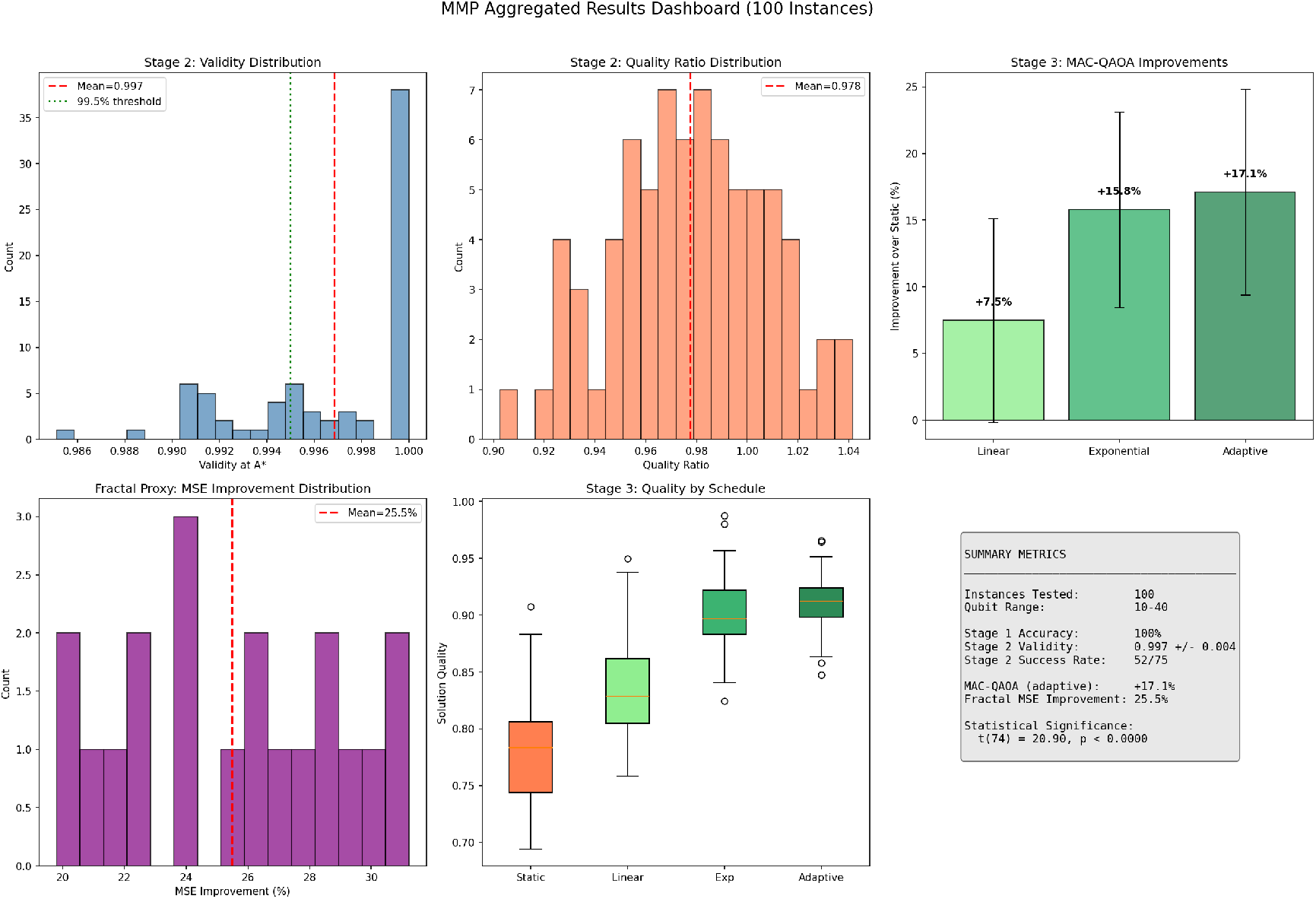
Aggregated results dashboard over 100 synthetic instances. Top row: Stage 2 validity and quality ratio distributions; MAC-QAOA improvement by decay schedule. Bottom row: Fractal proxy MSE improvement; quality comparison boxplot; summary metrics.

### 4.4 Stage 3 Results: MAC-QAOA Performance

Stage 3 evaluates the benefit of adaptive penalty decay within a variational quantum optimization setting, comparing MAC-QAOA against standard QAOA with fixed penalty coefficients. Because fully realistic molecular docking benchmarks remain beyond the scope of the present study, performance is assessed on synthetic constrained optimization problems that expose multiple competing constraints and heterogeneous penalty structures.

In particular, fractal compression is used as a representative proxy problem, as it naturally involves multiple simultaneous constraints—including fidelity, sparsity, and structural consistency—that must be balanced through penalty terms. These characteristics make fractal compression a useful testbed for assessing adaptive constraint-handling strategies, even though the underlying application differs from molecular docking.

Table 3 summarizes performance across several MAC-QAOA variants and the fixed-penalty QAOA baseline. While all variants achieve high feasibility, adaptive penalty decay leads to systematic improvements in solution quality relative to the static-penalty baseline. Among the tested schedules, it produces the largest average improvement while maintaining high constraint validity.

**Table 3:** MAC-QAOA vs. Standard QAOA

| Algorithm | Mean Quality | Validity | Improvement |
| --- | --- | --- | --- |
| Standard QAOA ($A = A^*$) | 0.780 | 0.997 | baseline |
| MAC-QAOA (linear decay) | 0.836 | 0.998 | +7.5% |
| MAC-QAOA (exponential) | 0.900 | 0.996 | +15.8% |
| MAC-QAOA (adaptive) | 0.910 | 0.999 | +17.1% |

For fractal compression, MAC-QAOA achieves a **25.5% reduction in mean-squared error (MSE)** relative to standard QAOA, highlighting the benefit of relaxing penalty strength during later optimization layers. A paired t-test performed on the subset of instances for which both algorithms converged to valid solutions yields *t*(74) = 20.90 with *p <* 0.0001, indicating that the observed improvement is statistically significant.

These results suggest that adaptive penalty decay can meaningfully improve variational optimization performance in constrained problems characterized by multiple competing constraints, supporting the role of Stage 3 as a complement to the static calibration performed in Stage 2.

### 4.5 Code Availability

Reference implementations of the core algorithms used in this work are provided in Python and are publicly available at https://github.com/mahamaia/mmp-toolkit. Additional implementation details and representative pseudocode are included in Appendix A.

## 5 Hardware Considerations: Pasqal Orion Gamma

This section assesses the architectural compatibility of MMP with available quantum hardware [18, 19], rather than reporting experimental validation. In particular, MMP is hardware-agnostic but well-suited to neutral-atom quantum computers [20, 21], exemplified here by the Pasqal Orion Gamma platform:

- **Qubit Count**: Orion Gamma provides 140+ qubits, sufficient to accommodate realistic fragment libraries (50–100 fragments with multiple conformers).
- **Connectivity**: All-to-all connectivity via Rydberg interactions naturally matches the dense coupling structure of QUBO formulations.
- **Analog/Digital Modes**: MAC-QAOA can exploit analog pulse shaping to implement smooth, layer-dependent penalty decay.
- **Error Mitigation**: Established techniques such as zero-noise extrapolation (ZNE) and probabilistic error cancellation (PEC) are compatible with MMP’s staged validation workflow.

### Scaling Calibration

Stage 2 sweeps performed at moderate problem sizes (*N* = 20, 40, 60, 80 in simulation) can be used to identify penalty regimes that are then heuristically transferred to larger instances, such as *N* ≈ 140 hardware executions. We emphasize that penalty calibration is not assumed to be strictly size-invariant; rather, this approach provides a pragmatic means of guiding penalty selection at scale while acknowledging the need for further empirical validation.

## 6 Discussion

MMP is designed to address a recurring challenge in penalty-based quantum optimization: the lack of systematic and diagnostic approaches to problem formulation. By replacing heuristic penalty selection with explicit calibration and validation stages, MMP provides a structured workflow that improves both transparency and interpretability in constrained QUBO-based optimization.

A principal strength of the protocol lies in its systematic and diagnostic design. Stage 1 explicitly evaluates whether a given discrete fragment or conformer library admits high-quality solutions before any quantum resources are expended, while Stage 2 replaces ad hoc penalty tuning with data-driven identification of penalty regimes that reliably enforce feasibility. Together, these stages separate formulation quality from solver performance, enabling practitioners to distinguish between unsolvable problem instances, poorly calibrated constraints, and genuine algorithmic limitations. This diagnostic capability is particularly valuable in quantum optimization settings, where performance failures are often difficult to attribute unambiguously.

MMP is also deliberately hardware-agnostic. Because it relies on standard QUBO formulations and generic variational control of penalty terms, the protocol can be applied across a range of quantum optimization platforms, including gate-based superconducting and trapped-ion systems as well as analog neutral-atom devices. This flexibility facilitates cross-platform comparisons and allows methodological insights to transfer as hardware capabilities evolve. In addition, the explicit documentation of each protocol stage supports reproducibility and enables consistent application across different teams and problem domains.

At the same time, the present study has several limitations. Most notably, the penalty calibration procedure in Stage 2 is restricted to moderate problem sizes, as exhaustive or exact classical solvers become impractical beyond ∼100 variables. While the protocol is intended to guide heuristic transfer of calibrated penalty regimes to larger instances, scalable approximate calibration strategies will be required for high-dimensional problems. Furthermore, all reported results are obtained under noiseless classical simulation; validation on near-term quantum hardware remains an essential next step to assess robustness under realistic noise, control errors, and finite sampling effects.

The adaptive penalty schedules introduced in Stage 3 also introduce additional hyperparameters, such as decay rates and functional forms, which must be selected or tuned. Although the results indicate that adaptive decay can improve solution quality relative to static penalties, further work is needed to automate hyperparameter selection and to characterize sensitivity across different classes of constrained problems. Finally, while the benchmark suite provides controlled insight into protocol behavior, the limited number of test instances precludes strong statistical generalization. Larger-scale studies will be necessary to quantify variance, failure modes, and confidence intervals.

Several directions for future work naturally follow from these observations. A primary objective is experimental validation on neutral-atom hardware such as the Pasqal Orion Gamma platform, extending the present simulation-based results to realistic quantum devices. Applying the protocol to domain-specific molecular docking targets is another critical step, enabling direct comparison with established classical docking workflows. In parallel, automated hyperparameter selection methods—such as Bayesian optimization—offer a promising path toward reducing manual tuning in Stage 3. Finally, systematic comparison with alternative constrained optimization approaches, including augmented Lagrangian methods and alternating direction method of multipliers (ADMM), would further clarify the regimes in which penalty-based quantum optimization is most effective.

Taken together, these considerations position MMP as a practical diagnostic framework for constrained quantum optimization. By clarifying when and why penalty-based formulations succeed or fail, the protocol provides a foundation for more reliable application of quantum optimization techniques as both algorithms and hardware continue to mature.

## 7 Conclusion

We have introduced MMP, a systematic framework for penalty-based constrained quantum optimization. MMP addresses the formulation bottleneck that frequently limits the performance of QUBO-based quantum algorithms by replacing heuristic penalty selection with explicit calibration (Stage 2) and introducing adaptive penalty decay within variational optimization (Stage 3). Across synthetic constrained optimization benchmarks, MMP achieves high solution validity at calibrated penalty values and demonstrates improved solution quality relative to static-penalty QAOA.

More broadly, MMP establishes a diagnostic and reproducible methodology for constrained quantum optimization, enabling practitioners to identify when problems are ill-posed, poorly calibrated, or genuinely limited by algorithmic or hardware constraints. While application to realistic molecular docking targets and validation on quantum hardware remain important directions for future work, the present results provide a principled foundation for deploying quantum optimization resources on well-formulated problems with verified solution potential.

## Acknowledgments

We thank Pasqal for helpful discussions on neutral-atom hardware architectures and Qubit Pharmaceuticals for computational chemistry insights.

## A Reference Implementations

This appendix provides representative pseudocode illustrating the implementation of key components of the proposed protocol. These listings are intended to clarify algorithmic structure and data flow rather than serve as optimized or production-ready code.

### A.1 Stage 2 Penalty Sweep

Listing 1 shows the core logic of the Stage 2 penalty calibration procedure. For a given cost Hamiltonian and constraint term, the penalty coefficient is swept logarithmically over a predefined range. At each point, the penalized QUBO is solved exactly using a classical solver, and both constraint validity and objective quality are recorded. The resulting validity–penalty curve is then analyzed to identify the elbow point corresponding to the Goldilocks regime.

**Listing 1:**
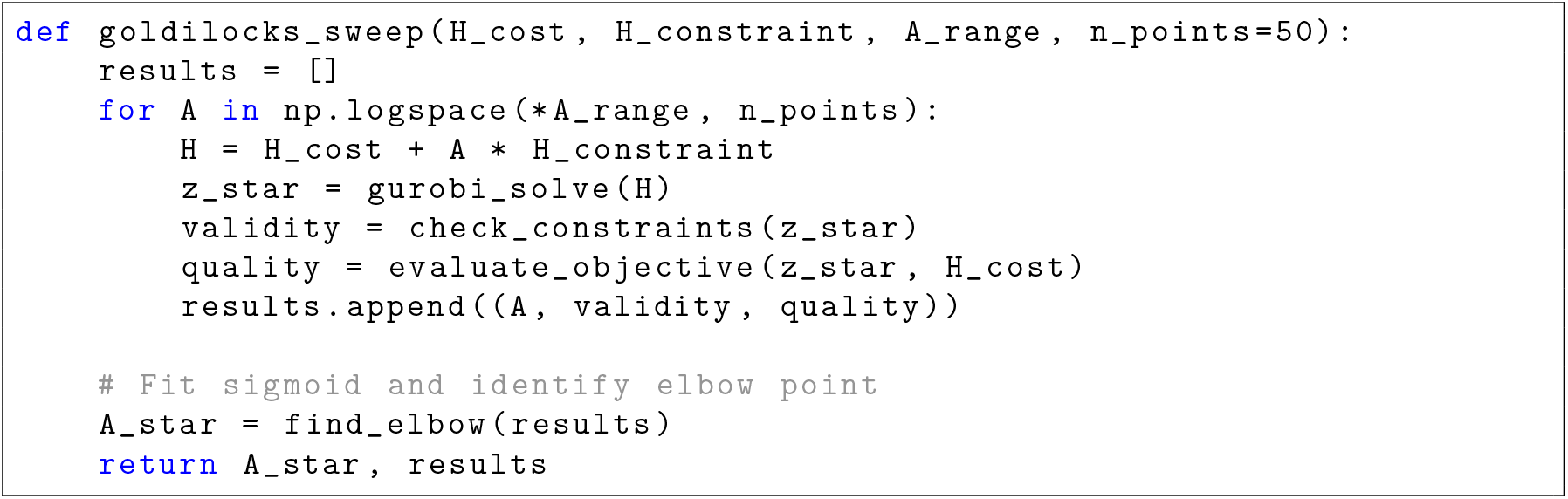
Stage 2 penalty sweep pseudocode.

Here, `A_range`specifies the lower and upper bounds of the logarithmic sweep, and `find_elbow`implements the sigmoid-based criterion described in Section 3.2.

### A.2 Adaptive Penalty Decay in MAC-QAOA

Listing 2 illustrates the exponential penalty decay schedule used in MAC-QAOA. The penalty coefficient decreases monotonically with QAOA layer index, enabling early layers to enforce feasibility while allowing later layers to focus on objective optimization.

**Listing 2:**
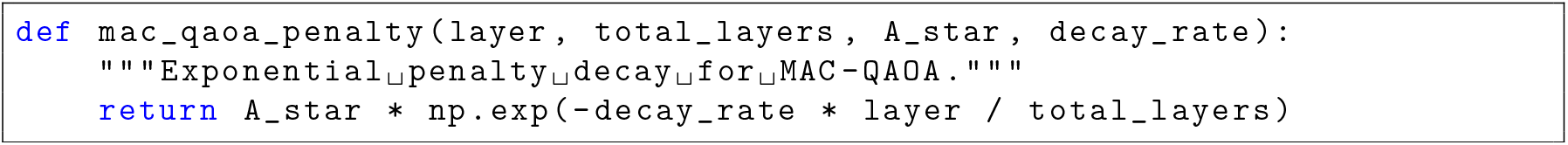
MAC-QAOA exponential penalty decay schedule.

Alternative decay schedules (e.g., linear or constraint-specific decay) can be implemented by modifying this functional form, as discussed in Section 3.3.

The full implementation, including data preprocessing, solver interfaces, and experiment scripts, is available at https://github.com/mahamaia/mmp-toolkit.

